# Transient protein-protein interactions perturb *E.coli* metabolome and cause gene dosage toxicity

**DOI:** 10.1101/066373

**Authors:** Sanchari Bhattacharyya, Shimon Bershtein, Jin Yan, Tijda Argun, Amy I. Gilson, Sunia Trauger, Eugene I. Shakhnovich

## Abstract

Several genes exhibit gene dosage toxicity yet its molecular underpinnings remain unknown. Here we demonstrate that overexpression of DHFR in *E. coli* causes toxic metabolic imbalance triggered by interactions with several enzymes involved in 1-carbon metabolism, in particular GlyA and PurH. DHFR overexpression partially inhibits activity of these enzymes, but at physiological concentrations, PurH-DHFR interaction enhances catalytic efficiency of DHFR, implying a functional interaction *in vivo*. Surprisingly, overexpression of orthologous DHFRs from other bacterial species caused minimal metabolic and fitness perturbations, despite pulling out more interacting partners than overexpressed endogenous DHFR. Orthologous DHFRs were less potent in inhibiting *E. coli* GlyA and PurH, or gaining a catalytic improvement upon interaction with PurH, indicating a partial loss of interaction specificity due to evolutionary divergence. This study shows how protein overexpression perturbs a dynamic network of weak yet potentially functional PPI with consequences for the metabolic state of cells and their fitness.

## Introduction

The experimental approaches to map the functional relationships between genes and phenotypes traditionally use perturbations of gene dosage via systematic deletion (Baba et al., 2006; Giaever et al., 2002; Pan et al., 2004; Sopko et al., 2006), down-regulation (Mnaimneh et al., 2004), or overexpression (Gelperin et al., 2005; Kitagawa et al., 2005; Makanae et al., 2013; Sopko et al., 2006) of the target genes. Phenotypes produced by gene overexpression generate a plethora of fitness effects that tend to deviate from those observed in gene deletion studies (Prelich, 2012; Sopko et al., 2006). High-throughput studies revealed that overexpression of a substantial fraction of genes is detrimental to fitness. In *E. coli*, overexpression of the majority of proteins is mildly to severely toxic under the conditions of the experiment (Kitagawa et al., 2005), whereas in yeast around 15% of overexpressed proteins produce morphological changes and drop in growth (Sopko et al., 2006). These findings are intriguing, given that gene-dosage increase plays a central role in evolutionary adaptations (Kondrashov, 2012), such as adaptations to a different carbon source (Brown et al., 1998), temperature (Riehle et al., 2001), and emergence of antibiotic resistance (Andersson and Hughes, 2009).

Several mechanisms were proposed to explain gene dosage toxicity (GDT), including resource overload (Makanae et al., 2013; Shachrai et al., 2010; Snoep et al., 1995; Stoebel et al., 2008), aggregation toxicity (Geiler-Samerotte et al., 2011; Kaiser et al., 2013), stoichiometric imbalance (Papp et al., 2003; Veitia et al., 2008), and non-specific PPIs (Ma et al., 2010; Vavouri et al., 2009). Vavouri *et al* hypothesized that GDT in yeast is predominantly caused by disordered proteins because of their potential involvement in multiple protein-protein and protein-DNA interactions, which, under the overexpression regime, could bring about deleterious mis-interactions (Vavouri et al., 2009). Sing and Dash hypothesized that electrostatic mis-interactions might be responsible for GDT in bacteria, which, unlike yeast, mostly lack proteins with disordered regions (Singh and Dash, 2013). Theoretical analyses suggested that the balance between functional and non-functional protein-protein interactions (PPI) is an important determinant of protein abundances in the cell (Deeds et al., 2007; Heo et al., 2011; Wallace and Drummond, 2015; Zhang et al., 2008). Thus, it is plausible that GDT might be caused by protein mis-interactions. However, specific mechanisms by which overexpression-induced protein mis-interactions cause toxicity and loss of fitness are not known.

Here we elucidate a molecular mechanism of GDT by focusing on molecular, systems, and organismal effects of overexpression of *E. coli* dihydrofolate reductase (EcDHFR). DHFR is an essential enzyme that catalyzes electron transfer reaction to form tetrahydrofolate, a carrier of single-carbon functional groups utilized in specific biochemical reactions (Harvey and Dev, 1975; Schnell et al., 2004). We explored fitness of *E. coli* in a broad range of EcDHFR abundances – from strong down-regulation to ~850 fold overexpression and established that the basal expression level is close to the optimal at which fitness is the highest, while *both* down regulation and overexpression appear toxic. While the drop in *E. coli* fitness upon DHFR downregulation is predicted by the enzymatic flux kinetics analysis (Bershtein et al., 2015; Rodrigues et al., 2016), toxicity upon overexpression is in variance with the prediction of this theory that predicts fitness neutrality once the functional capacity of an enzyme, defined as the product of its intracellular abundance and catalytic efficiency (*k_cat_*/*K_m_*), exceeds the maximal flux through a metabolic path (Dykhuizen et al., 1987; Feist and Palsson, 2010; Kacser and Burns, 1981; Lewis et al., 2012).

In order to get a detailed insight into the mechanism of GDT, we systematically analyzed perturbations in protein-protein interaction (PPI) and metabolic networks caused by overexpression of EcDHFR and its bacterial orthologues. We found that metabolic and fitness effects of overexpression were triggered exclusively by the endogenous DHFR, while the effects of overexpression of DHFR orthologues were much weaker on all scales. Pull-down assay coupled to LC-MS/MS revealed that over-expressed DHFR from *E.coli* and other bacteria interact with several enzymes in DHFR functional vicinity (folate pathway/purine biosynthesis), and *in vitro* analysis confirmed the presence of these weak interactions. However, the mutual effects on activity of DHFR and its interacting partners were markedly different between EcDHFR and its orthologues. These findings illustrate functional significance and ensuing evolutionary selection of weak transient protein-protein interactions in the crowded cytoplasm.

## Results

### *E. coli* is highly sensitive to DHFR dosage variation

Tetrahydrofolate, the product of DHFR activity, is utilized in numerous one-carbon metabolism reactions, including *de novo* purine biosynthesis, dTTP formation, and methionine and glycine production (Harvey and Dev, 1975; Schnell et al., 2004). As expected, a drop in DHFR activity and/or abundance, induced by either genetic changes (Bershtein et al., 2013; Bershtein et al., 2012; Bershtein et al., 2015), or DHFR inhibition (Kwon et al., 2008; Rodrigues et al., 2016; Sangurdekar et al., 2011), results in reduced fitness. But what is the effect of an *increase* in DHFR dosage on fitness? To answer this question, we designed an experimental system capable of generating controlled variation in a broad range of intracellular DHFR abundance in *E. coli*, and measured the effect of these changes on growth (Fig. 1A). A controlled drop in DHFR production was achieved by introducing LacI binding sites (*lacO*) in the chromosomal regulatory region of *folA* gene in a genetically modified *E. coli* strain constitutively expressing LacI (see Methods and (Bershtein et al., 2013)). Dialing down IPTG concentration (LacI repressor) resulted in a gradual decrease (over 100 fold) of the intracellular DHFR abundance (Fig. 1A). A gradual increase in DHFR abundance (from ~5 to 850-fold) was achieved by transforming WT *E. coli* strain with a plasmid carrying *folA* gene under the control of arabinose inducible pBAD promoter (see Methods and Fig. 1A,B). Fig. 1A shows that both drop *and* increase in intracellular DHFR abundance result in severe reduction (up to 73%) in growth rate. Fitness as a function of DHFR abundance therefore shows an ‘optimum’ at the basal DHFR level. However, the shape of the DHFR abundance – fitness function appears to be very different between DHFR depletion and DHFR overexpression regimes (Fig.1A). A drop in in fitness upon DHFR depletion is well described by Michaelis-Menten like dependence predicted by metabolic flux kinetics (Dykhuizen et al., 1987; Kacser and Burns, 1981) and verified experimentally (Bershtein et al., 2013; Bershtein et al., 2015; Rodrigues et al., 2016). In contrast, the increase in the intracellular DHFR levels above the optimal (basal) level leads to a steady and almost linear decline in fitness across the entire measured range of DHFR abundances (Fig. 1A,B). The qualitative differences between DHFR depletion and overexpression regimes clearly indicate that different mechanisms are responsible for loss of fitness at higher and lower ends of DHFR abundance-fitness curve.

**Figure 1:**
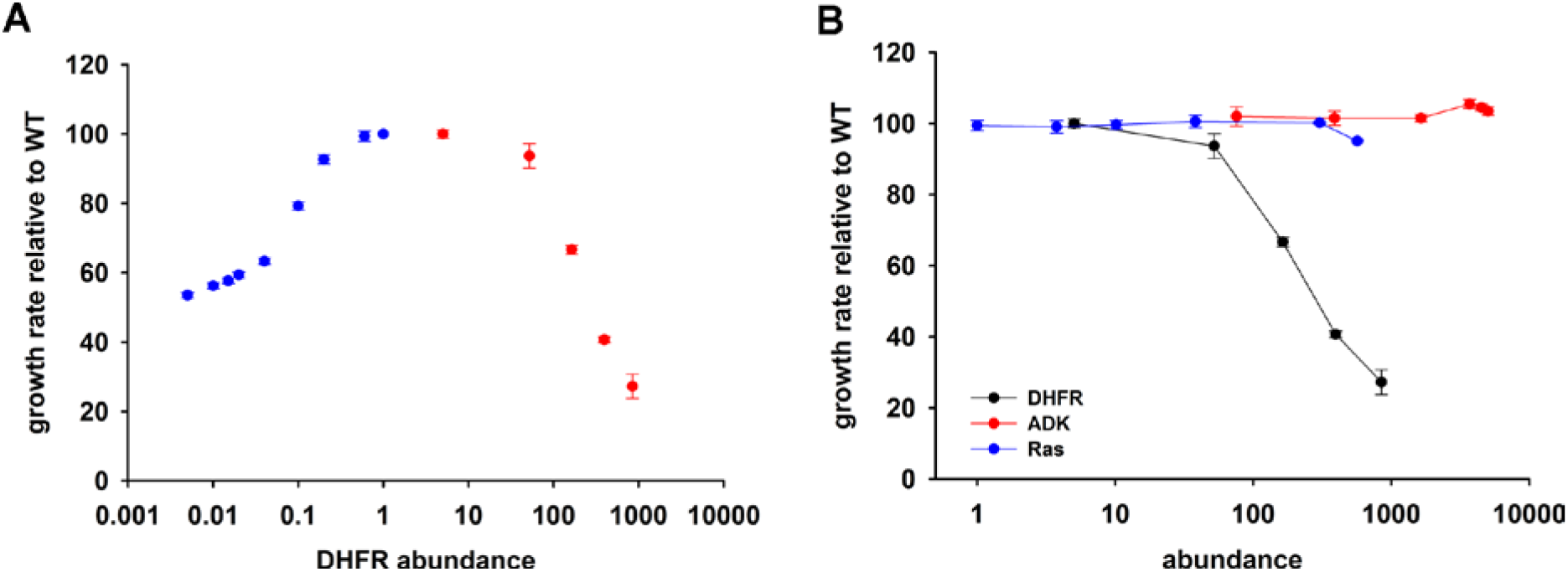
Over-expression of endogenous DHFR is detrimental to bacterial growth. (A) The effect of variation in DHFR dosage on fitness of *E. coli*. Change in the intracellular DHFR abundance (in log scale) is plotted against growth rate. DHFR abundance and bacterial growth are normalized against the parameters observed for wild-type strain. In case of over-expression, the growth rates were normalized with values obtained with no inducer (arabinose). Controlled drop (over 100 fold) in DHFR abundance (blue circles) was induced by IPTG titration of the chromosomal *folA* gene modified to contain LacI binding sites (see Methods and (Bershtein et al., 2013)). A controlled increase (~ 850 fold) in DHFR abundance (red circles) was achieved by arabinose titration of the wild-type strain transformed with a plasmid carrying the endogenous *folA* gene under the control pBAD promoter. The obtained abundance vs fitness function shows that the basal endogenous DHFR levels approach a physiological optimum with respect to *E. coli* growth rate. (B) Fitness as a function of protein abundance shown for EcDHFR, *E.coli* Adenylate Kinase (ADK) and a eukaryotic non-endogenous protein H-ras p21 (Ras). Only the expression of DHFR shows a dose-dependent toxicity and, therefore, it is not a generic effect of an over-expression burden. The intracellular DHFR, Ras, and ADK abundances were measured by Western Blot with custom raised antibodies (See Suppl. Materials and Methods). **See related** Fig.S1,S2

### DHFR dosage toxicity is not a result of resource overload or aggregation toxicity

Resource overload, or “protein burden”, is often invoked to explain the dosage toxicity of endogenous enzymes (Moriya, 2015), *e.g*. toxicity from lacZ production in the absence of lactose in *E.coli* (Dekel and Alon, 2005), and overexpression toxicity of glycolytic enzymes in yeast (Makanae et al., 2013) and *Z. mobilis* (Stoebel et al., 2008). To address the possibility that the observed toxicity of DHFR overexpression is a result of protein expression burden, we measured the effect of expression of two other control proteins on bacterial growth under identical conditions: an endogenous essential protein adenylate kinase (ADK), and a non-endogenous human H-ras p21 (Ras), - a small globular protein with size and fold similar to DHFR (de Vos et al., 1988) (Fig. 1B) At highest inducer concentration, Ras reached the abundance that is ~600 fold greater than of the endogenous DHFR, while ADK copy number reached ~5,000 fold increase over the physiological DHFR levels without causing any substantial drop in growth. Further, no toxicity was observed from an empty pBAD plasmid (Fig S1). Since *E. coli* DHFR and the control proteins are produced to comparable levels and from the same pBAD expression system, we conclude that the observed GDT of DHFR is protein specific and cannot be explained by either energetic or competition components of resource overload.

Fitness cost of overexpression is often attributed to aggregation of an overexpressed protein (Geiler-Samerotte et al., 2011). To test for possible aggregation of DHFR *in vivo, folA* gene in pBAD plasmid was fused in frame with the GFP coding gene and cellular fluorescence was analyzed. All overexpressing cells presented a highly diffuse fluorescence pattern indicative of lack of aggregation (Fig. S2A). A Western Blot analysis was carried out with the *E.coli* cell lysate containing pBAD-DHFR-GFP to confirm that the observed fluorescence emanated from the DHFR-GFP fusion protein and from GFP alone due to a possible proteolysis around the linker region of the fusion protein (Fig S2B, C). Thus, aggregation toxicity cannot cause GDT of DHFR overexpression.

### Overexpression of *E. coli* DHFR triggers a metabolic imbalance, while overexpression of DHFR bacterial orthologues does not

A subset of metabolic enzymes involved in more than one reaction can cause a bottleneck effect in some of the metabolic branches upon overexpression (Wagner et al., 2013). Although DHFR is a single-reaction enzyme, we tested whether its overexpression toxicity can be explained by a metabolic cost. To this end, we performed a metabolomics analysis of *E. coli* cultures overexpressing endogenous DHFR and ADK as a negative control. Whereas no measurable perturbation in metabolite levels could be detected upon ADK overexpression, a strong shift in metabolite pools was observed with overexpressed DHFR (Fig. 2A and Table S1). Specifically, we found a pronounced increase in AICAR and dUMP levels, accompanied by a drop in the levels of purines and pyrimidines (dTMP, dTTP, AMP, GMP, IMP). Toxicity could be partially rescued by adding these depleted metabolites (IMP, dTMP) to the growth medium (Fig. 2B and S3).

**Figure 2:**
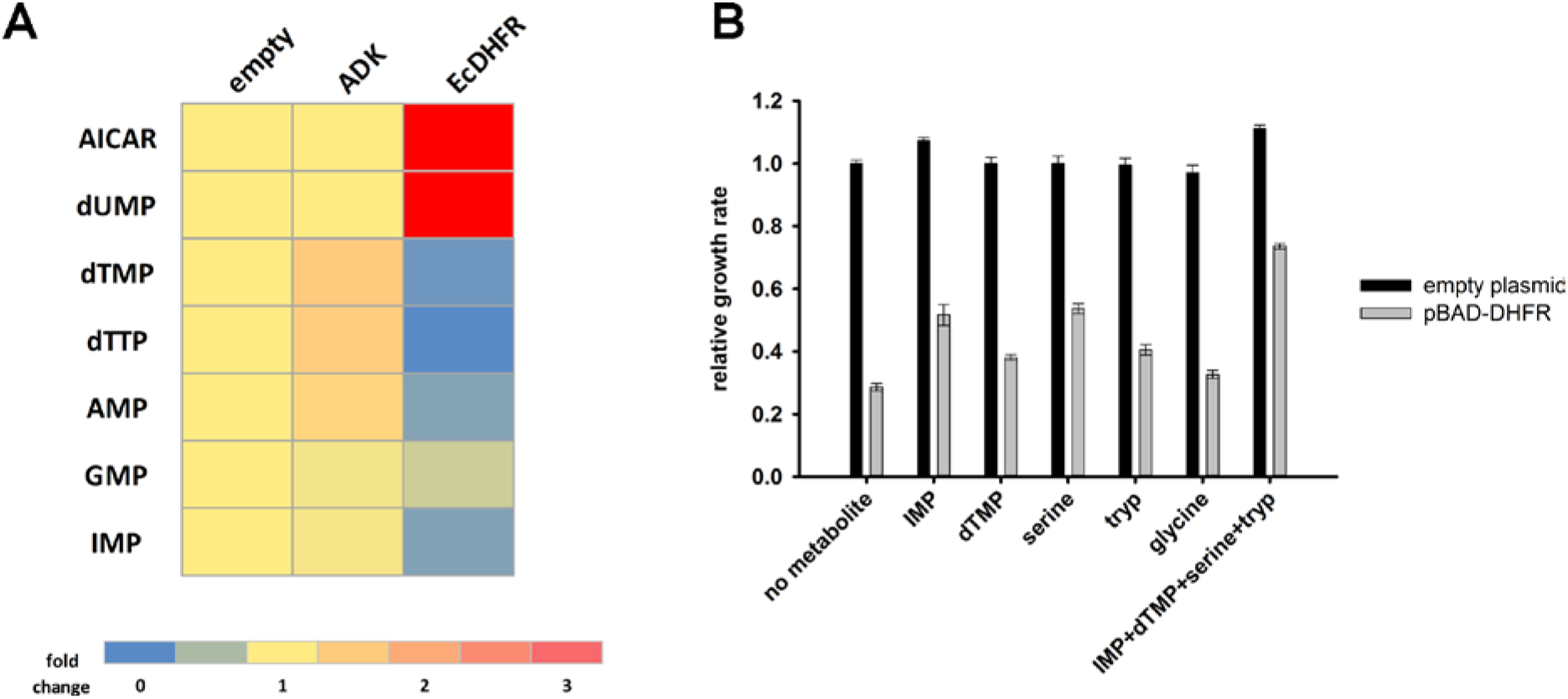
Overexpression of endogenous DHFR triggers a major metabolic shift. (A) Heat-map of the intracellular levels of various metabolites detected in bacterial cells over-expressing endogenous ADK and DHFR proteins from a pBAD plasmid. While ADK over-expression does not alter the metabolite levels, DHFR over-expression results in a pronounced up-regulation of AICAR and dUMP levels, and down-regulation of purines and pyrimidine nucleotides (dTMP, dTTP, AMP, GMP, IMP). (B) Metabolic complementation by addition of purine and pyrimidine nucleotides along with several amino acids, like serine and tryptophan, results in a partial rescue of fitness. 1mM of each of the metabolites was added to supplemented M9 medium at 37°C. Growth rates under different conditions were normalized by the growth rate obtained with empty plasmid without addition of any metabolite, and was referred to as ‘relative growth rate’. For all experiments the arabinose concentration used was 0.05 %, which corresponds to ~850-fold over-expression of DHFR and ~5000 fold increase of ADK. **See related** Fig.S3 **and** Table S1

Next, we tested the effect of overexpression of six distantly related but highly active DHFR orthologues from several bacterial species (Fig.S4 and Table S2). Remarkably, we found that these DHFRs had only a marginal effect on *E. coli* fitness (Figs. 3A,B). Furthermore, we found that over-expression of DHFRs originating from *Listeria innocua* (DHFR6) and *Bordetella avium* (DHFR11) did not result in any substantial change in metabolite levels. A small drop in IMP levels was observed for both, however AICAR levels were unperturbed (Fig.3C). Moreover, growth rate of all strains were highly correlated with their intracellular dTTP levels, indicating that depletion of metabolite levels is a direct signature of toxicity (Fig 3D). In summary, loss of fitness appears to be directly related to metabolite imbalance although excessive DHFR activity is not the root cause of this imbalance.

**Figure 3:**
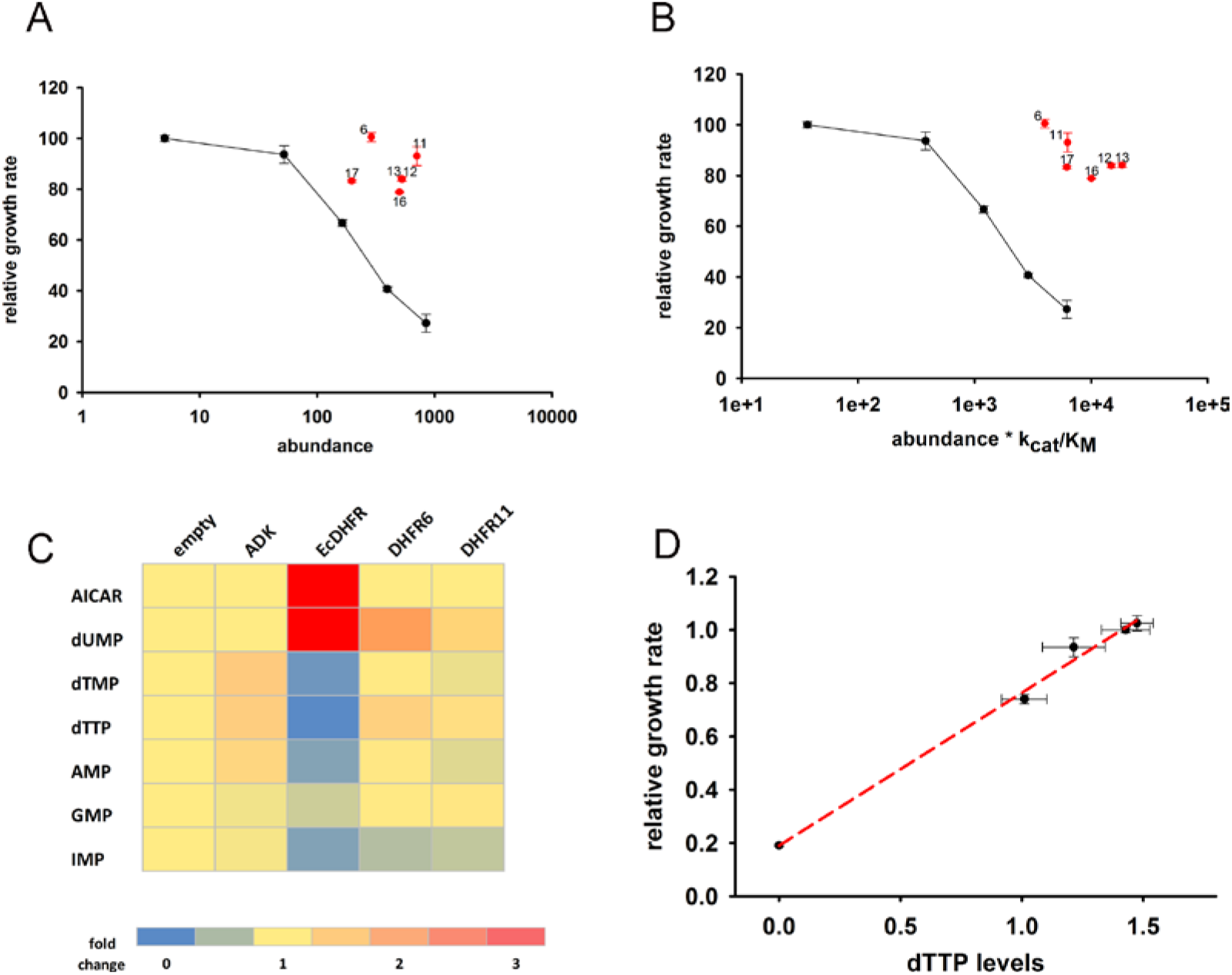
Orthologous DHFR proteins do not cause metabolic imbalance and dosage-related toxicity. Fitness is shown as a function of (A) intracellular DHFR abundance and (B) (abundance * k_cat_/K_M_) for overexpression of *E.coli* DHFRs as well as those from five mesophilic bacteria *Listeria innocua* (DHFR 6), *Bordetella avium* (DHFR 11), *Leuconostoc mesenteroides* (DHFR 12), *Aeromonas hydrophila* (DHFR 13), *Clostridium cellulolyticum* (DHFR 16) and *Streptococcus dysgalactiae* (DHFR 17) expressed from the same plasmid under pBAD-promoter. At equivalent concentration of functional DHFR proteins, only endogenous EcDHFR is highly toxic to *E.coli*. The intracellular abundance of orthologous DHFR was measured by Western Blot with anti-His antibodies (See Suppl. Materials and Methods). (C) Heat-map of the intracellular levels of various metabolites detected in cells overexpressing endogenous ADK and *E. coli* DHFR as well as two orthologous DHFRs (6 and 11). An empty pBAD plasmid is used as a control. Unlike *E.coli* DHFR, ADK and the orthologous DHFRs do not cause any major perturbation in the intracellular purine and pyrimidine levels (D) Growth rate of *E.coli* over-expressing ADK, *E.coli* DHFR, and orthologous DHFRs 6, 11 and 16 in the presence of 0.05% arabinose is shown as a function of the intracellular dTTP levels. All data were normalized by growth rate of *E.coli* transformed with the empty plasmid. The data shows that fitness is tightly correlated to cellular metabolite levels. **See related** Figure S4, Table S1, S2.

### Interacting partners of endogenous and orthologous DHFRs are enriched in 1-carbon metabolism enzymes

An increase in the intracellular concentration of proteins can induce potentially detrimental non-specific “mis-interactions” that are not present at the level of endogenous abundances (Heo et al., 2011; Maslov and Ispolatov, 2007; Vavouri et al., 2009; Zhang et al., 2008). Importantly, *E. coli* DHFR is a low copy monomeric enzyme (Taniguchi et al., 2010) with no physical interactions detected with other proteins when used as a prey at *its basal level (Hu et al., 2009). Further, no DHFR interactions were detected in yeast*, and only 1 in Drosophila [dip.doe-mbi.ucla.edu]. We sought to determine possible interacting partners of *over-expressed* EcDHFR and two controls, ADK and Ras. To that end protein complexes in cells were stabilized *in vivo* by a cell-permeable cross-linker, followed by cell lysis, immunoprecipitation (IP) performed on soluble lysates, and LC-MS/MS analysis (See Fig. 4A and Methods for details). As shown in Table S3 and S4, overexpressed DHFR pulled out several interaction partners, while ADK and Ras picked very few (3-4), suggesting that cellular PPI is the most likely reason for DHFR overexpression toxicity. To compare the PPI profiles of *E.coli* DHFR vs the orthologues, a similar IP LC-MS/MS analysis was performed with the exception that mouse monoclonal anti-His antibodies were used, as opposed to polyclonal rabbit anti-EcDHFR, ADK, and Ras antibodies (since orthologs do not bind anti-EcDHFR antibodies due to large difference in sequence identity). As a first step, we compared the PPI profiles of EcDHFR with two different antibodies and found that the anti-his antibody pulled out a smaller number of interacting proteins (Table S3). This is probably because cellular PPI that target regions close to the N-terminal His-tag of DHFR cause steric hindrance for binding of anti-His antibodies. Despite this caveat, we went ahead to compare the PPI profiles of EcDHFR and orthologous DHFR 11 and found that DHFR 11 pulled out many more proteins than EcDHFR (Table S3 and S4). Interestingly, almost all proteins constituting the EcDHFR interactome can be found in the DHFR11 interactome. Why is then overexpression toxicity limited only to EcDHFR? A closer look into the set of 29 interacting proteins of EcDHFR that were common to both anti-EcDHFR and anti-his antibody pull-downs (Sheet 1 of Table S4) allowed us to classify them according to their function (Sheet 2 of Table S4). While several proteins belong to carbohydrate metabolism and amino acid biosynthesis, there were several enzymes closely related to DHFR function, such as 1-carbon metabolism. Tetrahydrofolate, the product of DHFR is immediately taken up by Serine Hydroxymethyltransferase (GlyA) to produce 5,10-methylene-THF, which in a series of steps is subsequently used to produce dTMP from dUMP (Fig 5B). 5,10-methylene-THF is also utilized in a second pathway to produce ^10^Nf-THF, which is then consumed in the last step of purine biosynthesis by PurH. Hence, partial inhibition of either of these paths can explain the metabolic imbalance and fitness cost associated with DHFR overexpression.

**Figure 4:**
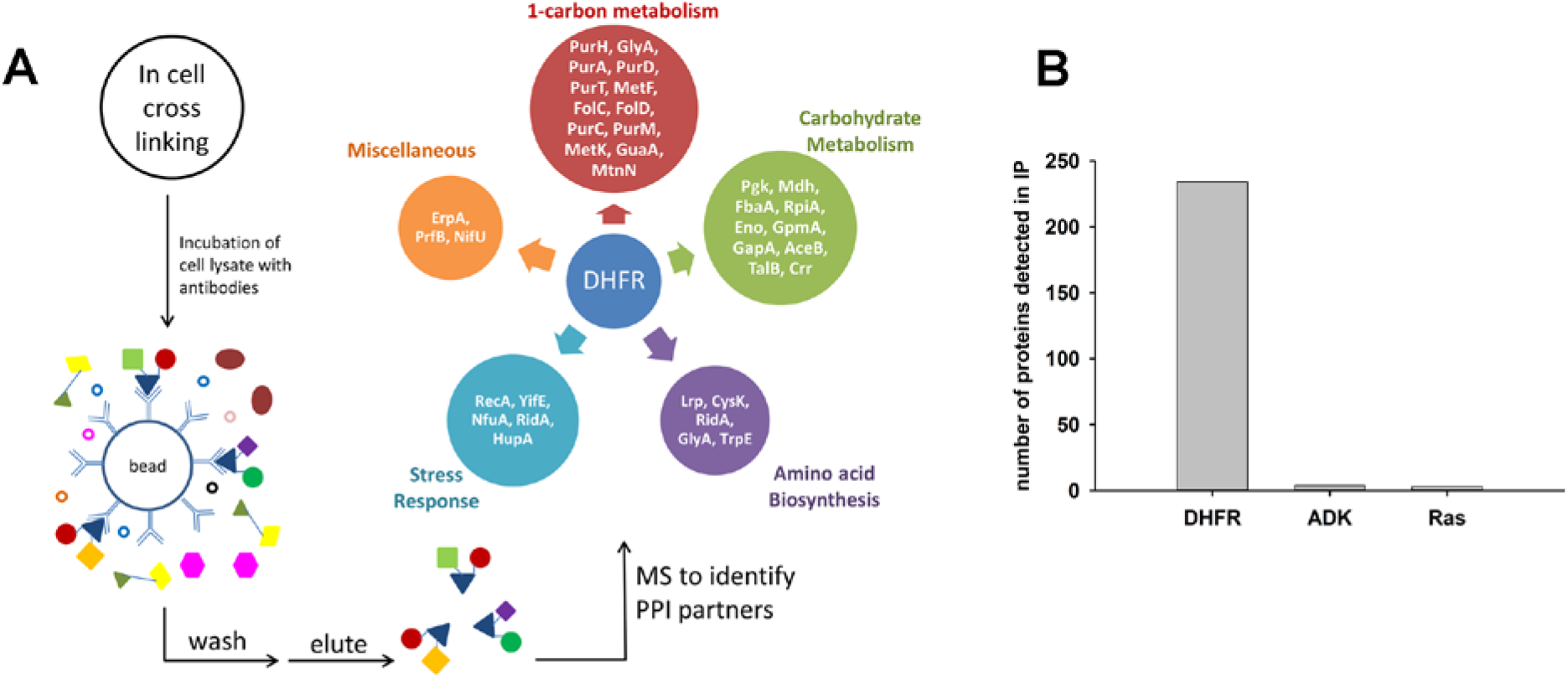
Detection of protein-protein interaction (PPI) partners of *E.coli* DHFR. (A) Schematics of the co-immunoprecipitation protocol. Proteins were cross-linked inside the cell using a cell-permeable cross-linker DSP (dithiobis(succinimidyl propionate)), and following lysis, the lysate was captured on anti-DHFR/anti-ADK/anti-Ras/anti-His-tag antibodies to fish out the protein of interest and its PPI partners. The interacting partners were subsequently identified using LC-MS/MS analysis. The observed interactome of overexpressed DHFR could be classified broadly into carbohydrate metabolism and 1-carbon metabolism (purine biosynthesis and amino acid metabolism groups). (B) Over-expressed *E.coli* DHFR picked out 234 proteins in total, while ADK and Ras detected only 4 and 3 proteins respectively. All IP datasets were background subtracted using lysate from *E.coli* cells that were transformed with empty pBAD plasmid. There was no overlap among the interactomes of DHFR, ADK and Ras, indicating that these were specific interactions. (**see related Tables S3 and S4**)

**Figure 5:**
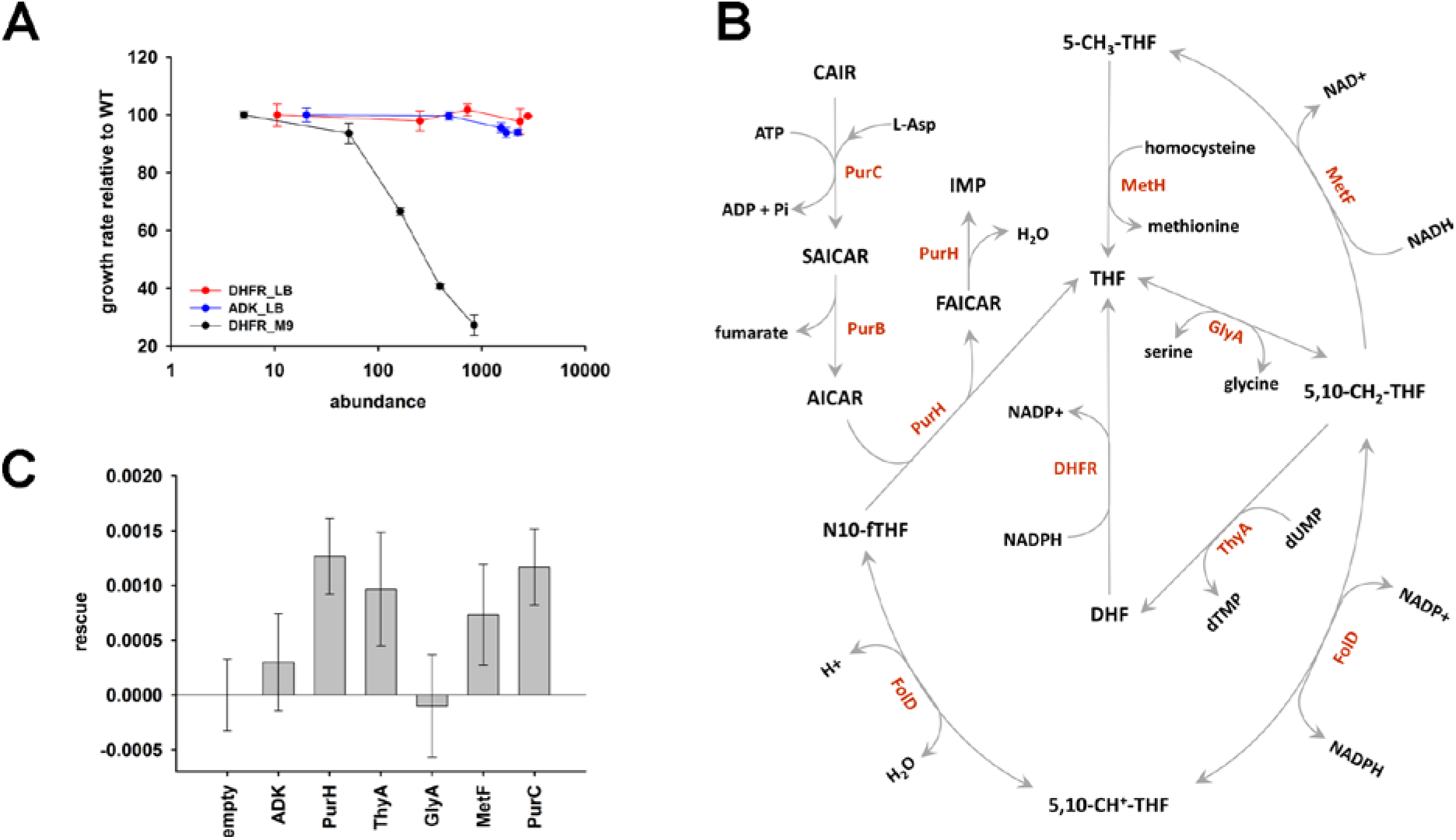
The role of purine biosynthesis and 1-carbon metabolism enzymes in DHFR over-expression toxicity. (A) Unlike supplemented M9 medium, over-expression of DHFR is not toxic in rich medium (LB) in terms of growth rate (note, however, that saturation ODs at 600nm is lower in LB in comparison to growth in M9; see Fig S5). This indicates the role of conditionally essential genes like *purH, purC, metF, etc*. in determining the over-expression toxicity of DHFR. All data were normalized by growth rate of *E.coli* transformed with the empty plasmid. (B) Schematics of the 1-carbon pathway metabolic pathway. GlyA, ThyA, MetF are important enzymes that function immediately downstream of DHFR and utilize THF or its derivative. (C) Partial rescue of fitness of DHFR over-expressing *E.coli* by a dual-expression system. DHFR was expressed from a IPTG-inducible plasmid while PurH, ThyA, GlyA, MetF and PurC along with a negative control protein ADK were expressed separately from an arabinose-inducible pBAD system. Rescue factor is defined as (*μ*(*DHFR* + *X*) − *μ*(*DHFR* + *empty*)) − (*μ*(*X*) − *μ*(*empty*)), where *μ* is the growth rate, and X is the corresponding protein **See related** Figs.S5 **and** S6

### DHFR overexpression toxicity is caused by PPI with essential enzymes

*De novo* purine biosynthesis enzymes are essential for growth in minimal media, such as M9, but are non-essential in LB medium that contains purines (Gerdes et al., 2003; Joyce et al., 2006). This observation suggests that if inhibition of purine biosynthesis through PPI with overexpressed DHFR is partly responsible for GDT, DHFR overexpression would be less toxic in LB. Indeed, we found that cells overexpressing DHFR do not show a reduction in “growth rates” when grown in LB (Fig. 5A), however there was a gradual drop in the saturation ODs at higher expression levels of DHFR (Fig S5A). This might be due to DHFR interaction with other 1-carbon metabolism enzymes (GlyA, ThyA) or enzymes belonging to carbohydrate metabolism, namely phosphoglycerate kinase, fructose-biphosphate aldolase, which are essential even in LB (Baba et al., 2006). Further, we ran several complementation experiments in which we tested the effect of overexpression of enzymes involved in 1-carbon metabolism (schematically depicted in Fig. 5B) on rescuing the DHFR overexpression toxicity. Most of these enzymes (with the exception of ThyA) were shown to interact with overexpressed DHFR in the IP/LC-MS/MS assay (Fig.4, Table S4). As shown in Fig. 5C, overexpression of PurH, PurC, ThyA, and MetF resulted in a moderate yet significant alleviation of DHFR dosage toxicity, validating some of the interactions detected in IP. However the rescue was only moderate, as most of these proteins themselves were toxic upon over-expression (Fig S6). A protein involved in tryptophan biosynthesis, TrpE, was detected in IP, which, if sequestered, might lead to a drop in tryptophan levels in the cell. Indeed addition of tryptophan to the growth medium partly rescued the growth defect of DHFR over-expression (Fig.2B and S3), again validating the functional significance of the IP-detected interactions.

### In vitro analysis of DHFR interactions with its metabolic neighbors

Analysis of 1-carbon metabolism pathway suggested that effect of DHFR on several potential interaction partner proteins could explain, at least partially, the significant metabolic shifts and fitness effects of DHFR overexpression. Of them, we chose GlyA and PurH proteins for in depth *in vitro* analysis. GlyA works immediately downstream of DHFR in the folate pathway and uses THF to catalyze the reversible interconversion between glycine and serine (Fig. 5B). Although complementation with overexpressed GlyA did not lead to partial rescue of fitness, presumably due to its intrinsic toxicity (Fig.S6), addition of serine did lead to a partial alleviation of toxicity of overexpressed EcDHFR (Fig.S3).

PurH is a bifunctional enzyme having IMP cyclohydrolase/aminoimidazole carboxamide ribonucleotide (AICAR) transformylase activities that catalyzes the last two steps in *de novo* purine synthesis (inosine-5’-phosphate biosynthesis path). A direct inhibition of both activities might explain the accumulation of AICAR and drop in IMP, AMP and GMP in the metabolite pools of overexpressing DHFR cells. Indeed, complementation with additional PurH protein showed a partial rescue of fitness (Fig. 5C).

As a first step, we purified GlyA and PurH from *E. coli* and measured their interaction with *E.coli* and orthologous DHFRs *in vitro* using surface plasmon resonance (SPR). Both proteins were found to interact weakly with DHFR (Fig.S7 and Table S5). GlyA interacts with EcDHFR with a K_D_ of 8μM and with DHFR6 and 11 with K_D_ of 5.3μM and 4μM respectively. PurH interacts with both EcDHFR and two orthologs (DHFR6 and DHFR11) with very similar K_D_ in the low μM range (Fig S7 and Table S5). ADK served as the negative control for all binding experiments and did not show any binding to GlyA and PurH. The similarity of the interaction between orthologous DHFRs and their interacting partners (GlyA and PurH) suggests that binding alone is insufficient to explain the observed selective toxicity of EcDHFR overexpression.

### E.coli DHFR affects activity of its interacting partners *in vitro*

The binding studies did not reveal significant differences in interaction of *E.coli* and orthologous DHFRs with GlyA and PurH. As a next step, we analyzed the *functional* effect of *E.coli* and orthologous DHFRs on the partner proteins. For GlyA, the formation of a ternary quinoid complex in the presence of pyridoxal phosphate (PLP), 5f-THF and glycine was used as a measure to determine its activity (Schirch et al., 1985) (see Methods). While presence of ADK did not alter the ability of GlyA towards complex formation, *E.coli* as well as the orthologous DHFRs did show a concentration dependent inhibition of GlyA (Fig 6A). However, EcDHFR is clearly a more potent inhibitor of GlyA than its orthologs (Fig 6B). In addition, binding of EcDHFR to GlyA resulted in the largest increase in K_D_ of 5f-THF binding (Fig S8A and B).

**Figure 6:**
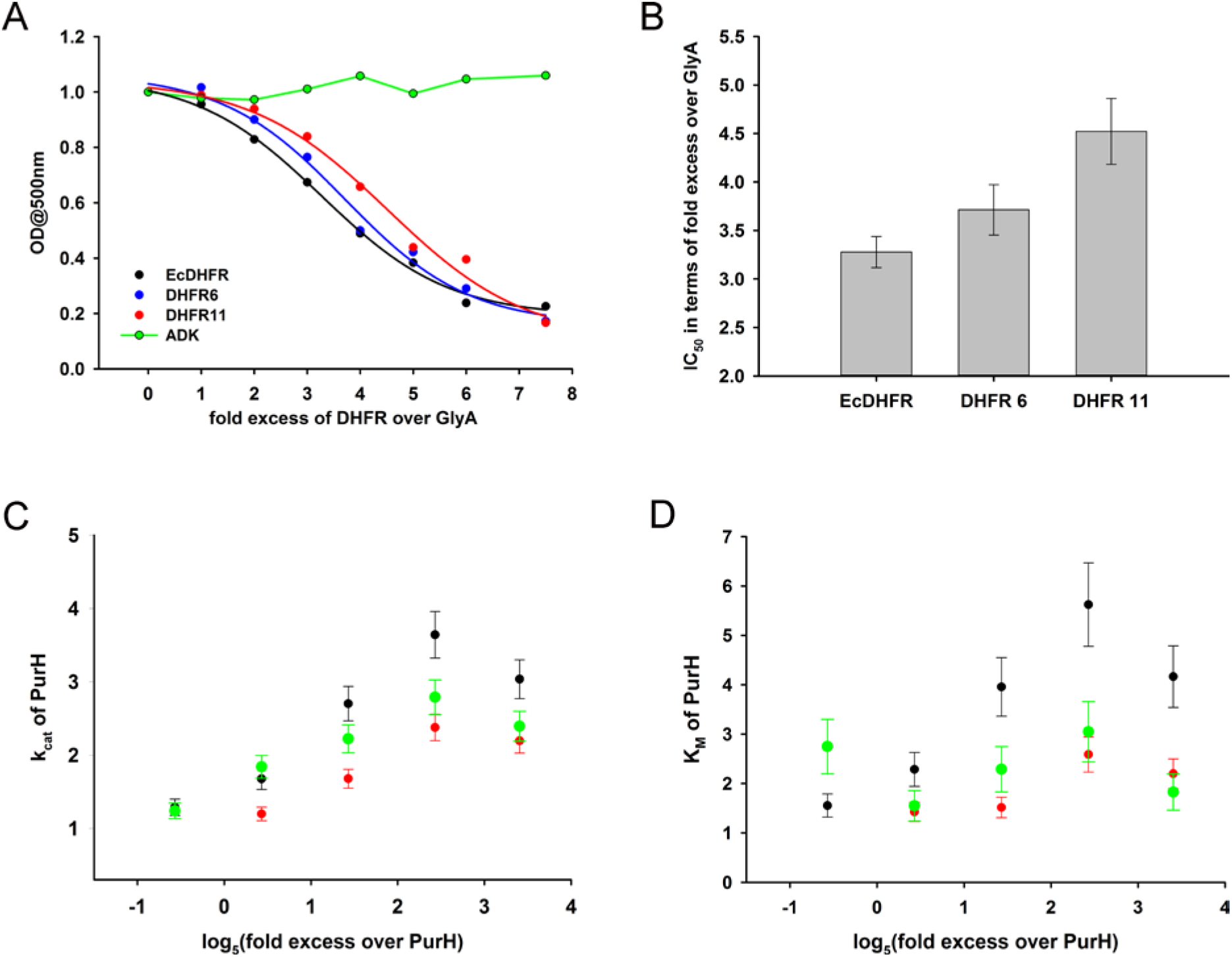
Compared to orthologous DHFRs, *E.coli* DHFR is a more potent inhibitor of GlyA and PurH. (A) Formation of a ternary complex (GlyA-PLP+glycine+5f-THF) was monitored at 500 nm as a function of increasing amounts of different DHFRs and the negative control protein ADK. 20μM GlyA was pre-incubated with varying concentrations of DHFRs and ADK (zero to 150μM) before 0.2M glycine and 200μM 5f-THF were added to it. All data were normalized by those in the presence of an equal volume of buffer. Data were fit to a 4-parameter sigmoidal function to extract the fold excess of DHFR required to achieve 50% inhibition (IC_50_) as shown in panel (B). Though all DHFRs caused substantial inhibition of GlyA compared to ADK, EcDHFR has the lowest IC_50_, explaining its higher potency in sequestering GlyA and, hence, toxicity. Catalytic rate *k_cat_* (C) and Michaelis coefficient *K_M_* (D) of PurH for ^10^Nf-THF measured in the presence of different DHFRs and ADK. All values were normalized relative to those of PurH measured in the absence of added protein. 250nM of PurH was pre-incubated with varying concentrations of DHFRs or ADK (0.1μM, 0.5μM, 2.5μM, 12.5μM and 60μM) and, subsequently, the initial rate of transfer of formyl group (AICAR transformylase activity) from ^10^N-formyl THF to AICAR was measured at 298nm. For determination of *k_cat_* and *K_M_* at each concentration of a protein, the concentration of ^10^Nf-THF was varied from 20μM to 1mM, while AICAR concentration was fixed at saturation (500μM). For all proteins, *k_cat_* increased with increasing concentration of protein added, dropping off slightly at the highest concentration. The mechanism of this effect is not fully understood, but can be partially attributed to the generic crowding effect of proteins. However, only EcDHFR caused a concentration dependent increase in *K_M_* of PurH for ^10^Nf-THF, thereby explaining its selective toxicity upon over-expression. **See related** Fig.S8,S9 **and** Table S5.

We next assayed the AICAR transformylase activity of PurH in the presence of *E.coli* and the orthologous DHFRs, and the negative control protein ADK. EcDHFR was found to have the strongest concentration-dependent effect on both ***k_cat_*** and ***K_M_*** of AICAR transformylase activity of PurH (Fig 6C, D and Fig S9) with especially strong and pronounced detrimental effect on ***K_M_***. In contrast, addition of the same amount of DHFR11 did not affect the activity of AICAR transformylase to a significant degree. In summary, despite the similarity in binding parameters, the functional outcome of the interaction of DHFR with metabolically related enzymes GlyA and PurH is strikingly different for EcDHFR compared to the orthologs, and might, therefore, be a cause of selective toxicity of EcDHFR.

### The effect of PurH on DHFR activity at physiological concentrations is beneficial

The data so far suggest that over-expressed DHFR interacts with several proteins, including GlyA and PurH and potentially compromises their function. Importantly, the effect is the strongest when DHFR and the interacting partner are from *E. coli*. This observation points out to an intriguing possibility that the interactions might have a functional significance even at the basal expression levels of DHFR, but are too transient to be detected. To test this assumption, we analyzed the mutual effect of PurH and DHFR on each other’s activity *in vitro* at physiological concentrations of both enzymes. While DHFR does not affect AICAR transformylase activity of PurH at physiological concentration (Fig.6C,D), we did observe a strong effect of PurH on activity of DHFR, which is highly specific to the *E. coli* variant (Fig. 7). Both PurH and ADK resulted in a moderate increase in *k_cat_* of all DHFRs (Fig 7A), however the presence of PurH led to a significant decrease in *K_M_* of EcDHFR, while it had almost no or weak opposite effect on *K_M_* of the orthologues (Fig.7B). As a result, at physiological concentrations PurH significantly increases the catalytic efficiency (*k_cat_* / *K_M_*) of EcDHFR, whereas the activity of its orthologues remains unaffected (Fig.7C).

**Figure 7:**
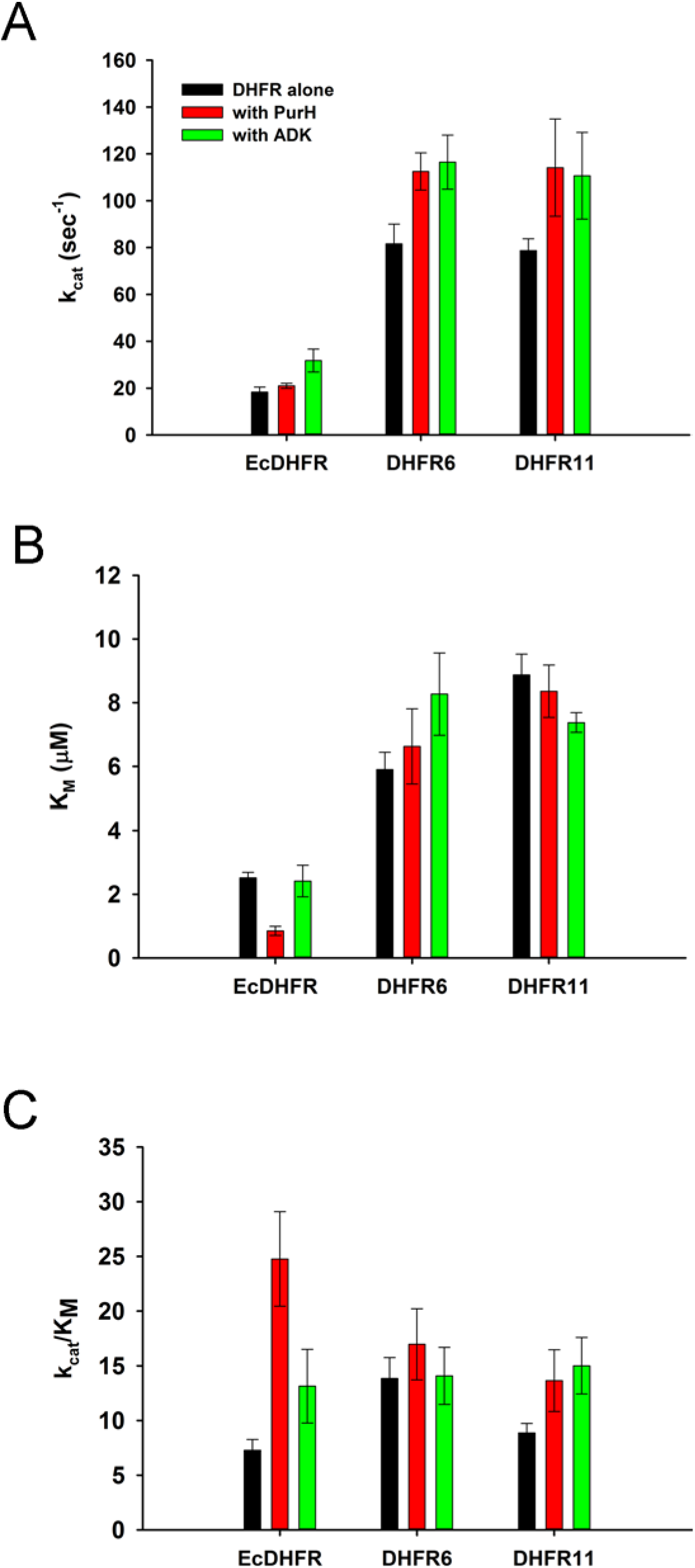
PurH has a selective beneficial effect on EcDHFR activity only. (A) Catalytic rate (*k_cat_*), Michaelis coefficient (*K_M_*), and the catalytic efficiency (*k_cat_*/*K_M_*) of EcDHFR, DHFR6 and DHFR11 for dihydrofolic acid (DHF) in the presence of low concentrations of PurH. 10nM DHFR were pre-incubated with 15nM of PurH or ADK and, subsequently, the initial rate of conversion of NADPH to NADP+ was measured at 340nm. For determination of *k_cat_* and *K_M_*, the concentration of DHF was varied from 0.1μM to 16μM for EcDHFR and from 1μM to 64μM for DHFR6 and 11, while NADPH concentration was fixed at saturation (150μM). Both PurH and ADK resulted in an increase in *k_cat_* of all the DHFRs, however PurH caused a significant drop in *K_M_* for EcDHFR only. Therefore, at physiological ratios of proteins, PurH is only beneficial for EcDHFR. This observation suggests an evolutionary functional relationship between PurH and EcDHFR at physiological concentrations. Orthologous DHFRs that have diverged during the course of evolution no longer have this benefit from *E.coli* PurH.

### Excess of DHFR converts transient interactions into permanent ones

Tetrahydrofolate, the product of DHFR activity is a precursor for 5,10-methylene THF, which is used for dTMP synthesis, as well as ^10^Nformyl-THF, which is eventually utilized by PurH in the last step of the *de novo* purine biosynthesis pathway to produce IMP. Hence, DHFR is functionally very closely connected to the family of purine/pyrimidine biosynthesis enzymes. Purine biosynthesis enzymes were found to be spatially localized forming the ‘purinosome’, in mammalian cells, but no evidence exists for *E. coli* (An et al., 2008; Deng et al., 2012; French et al., 2016). It is possible that at basal expression levels DHFR is a member of a dynamic ‘metabolon’ that facilitates channeling of tetrahydrofolate or its derivatives. However, when over-expressed, the toxicity is triggered by a stoichiometric imbalance of the complex which leads to conversion of these transient complexes to more permanent ones.

To assess the feasibility of such a scenario we use the Law of Mass Action to determine the fraction of DHFR and PurH in the monomeric form and in complexes:

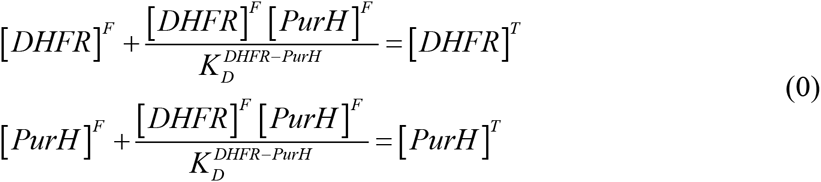

where square brackets denote the concentrations of enzymes, and superscripts *F* and T refer to free and total enzyme concentrations, respectively. 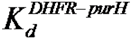 is the equilibrium binding constant between PurH and DHFR. Eq. 1 represents simple stoichiometry relations for both enzymes.

To solve Eq.1, we used 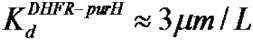 determined in this study (Table S5), and known endogenous cellular concentrations of DHFR and PurH of 5×10^−8^ and 2×10^−7^ moles/L respectively (corresponding to 50 and 200 molecules per cell (Taniguchi et al., 2010)). We found that at the endogenous abundance levels interaction between DHFR and PurH is transient with only 5% of DHFR found in complex with PurH and 1% of PurH is in complex with DHFR at any given time. However, when DHFR is ~1000 fold overexpressed, only 4% PurH remains free in cytoplasm, while the remaining 96% are complexed with DHFR. Similar estimates apply to GlyA.

## Discussion

High-throughput studies showed that GDT is a common phenomenon affecting thousands of genes (Kitagawa et al., 2005; Sopko et al., 2006). However, despite its importance, there are no experimental “case studies” that uncover molecular underpinnings of GDT. Here we carried out a study whereby we focus on a specific gene, *folA*, and delineate, in great detail and on multiple scales, the molecular mechanism of DHFR dosage toxicity in *E. coli*. We showed that fitness cost of DHFR overexpression could only be understood by integrating the metabolic and protein-protein interaction networks of the organism. Specifically, interactions between DHFR and several other metabolic enzymes, lead to a reduced flux through the purine, pyrimidine and amino acids synthesis paths. The imbalance in protein-protein interactions caused by overexpression of DHFR leads to a dead-end metabolic imbalance, such as the accumulation of AICAR – a precursor of inosine-5’-phosphate, and reduced growth. Overexpression of highly active but diverged orthologous DHFR caused only minor change in metabolite pools and fitness, indicating that neither protein burden nor metabolic cost *per se* can explain the endogenous DHFR dosage toxicity. The comprehensive IP/LC-MS/MS analysis confirmed by *in vitro* measurements of selected pairs of proteins clearly point out to spurious PPI formed between the overexpressed DHFR and several other metabolically and functionally related proteins in *E. coli* cytoplasm as the most probable cause of GDT.

The impact of overexpression on all levels of cell organization – molecular, systems, and phenotypic, dramatically differs between “self”, *i.e. E.coli*’s, and “foreign” DHFR. This key finding clearly suggests the crucial role of evolutionary selection in shaping PPI at the whole proteome level. Remarkably, we found that at the physiological concentrations weak transient PPI can enhance the enzymatic activity of DHFR, provided that the interacting proteins are *from the same organism*. Further, we found that overexpressed “foreign” DHFR pulls up many more interaction partners from *E. coli* proteome than does EcDHFR, suggesting that DHFR was subjected to selection against promiscuous mis-interactions in *E. coli* cytoplasm. Thus, evolution of the PPI network entails both positive *and* negative selection for transient interactions. Transient interactions between a select set of metabolically related proteins might be beneficial because they can enhance catalytic efficiency through ligand channeling. Conversely, they can be selected against to avoid massive non-functional promiscuous PPI. The weaker binding of EcDHFR to GlyA (Table S5) may also be indicative of a selection against transient interactions, as higher copy number enzymes, like GlyA, (expressed at ~10,000 copies/cell, (Taniguchi et al., 2010)) are more prone to mis-interact, comparatively to less abundant proteins, such as PurH (expressed at ~200 copies per cell (Taniguchi et al., 2010)). Earlier theoretical analyses envisioned both scenarios: selection against massive non-functional PPI (Heo et al., 2011; Levy et al., 2012; Vavouri et al., 2009; Yang et al., 2012; Zhang et al., 2008), and selection for interactions between metabolically related enzymes (Durek and Walther, 2008; Huthmacher et al., 2007, 2008). This work provides experimental evidence that both these factors are indeed at play in evolution of PPI and metabolic networks.

Several authors hypothesized that sequestration of functional proteins into non-productive promiscuous PPI upon overexpression may cause GDT (Heo et al., 2011; Vavouri et al., 2009; Yang et al., 2012). Our study also shows that GDT of DHFR is triggered by potentially weak functional interactions gone “awry” under the overexpression regime that causes a stoichiometric PPI imbalance and formation of permanent complexes in lieu of transient ones. However, unlike these studies, we show that the actual cause of the overexpression toxicity is the metabolic imbalance that ensues once the permanent PPIs are formed. Apparently, actual ‘mis-interactions’ are formed upon overexpression of orthologous DHFR proteins. In contrast to the previous studies, however, these interactions happen to be much less toxic than those picked by *E.coli* DHFR, at least at the experimental level of resolution. Nevertheless, weaker fitness effects of mis-interactions can affect evolution of the proteome to minimize promiscuous PPI (Heo et al., 2011; Levy et al., 2012), and our finding that non-endogenous DHFRs pulls out more interacting partners than EcDHFR points in this direction.

Broad distribution of protein abundances (Ghaemmaghami et al., 2003; Ishihama et al., 2008) and remarkable conservation of abundances of orthologous proteins between species (Laurent et al., 2010; Schrimpf et al., 2009) strongly suggests that protein abundance is a selectable trait. In contrast to protein stability, whose landscape is largely monotonic, at least for some proteins (Bershtein et al., 2012; Bloom et al., 2006; Rodrigues et al., 2016), fitness landscape for protein abundances is non-monotonic for many proteins, reaching an optimum at certain levels. While loss of fitness at low abundance is intuitively associated with loss of function, the GDT at high abundance is a system-level collective effect, related to interactions with multiple proteins as well as the link between PPI and metabolic networks, as shown here for DHFR. Apparently, specific functional demands dictate certain minimal amounts of proteins. Evolution of biophysical and biochemical parameters (activity, stability, and affinity to a functional protein and DNA partners) provide significant leverage to avoid prohibitively high expression levels which might cause GDT. Nevertheless, there is a certain evolutionary cost of maintaining high levels of activity and/or functional interaction strength due to supply of potentially deleterious mutations. Future studies will fully reveal the extent to which GDT affects evolved distributions of molecular properties of proteins and their abundances.

## Methods

### Metabolite extraction

BW27783 cells transformed with pBAD-empty, pBAD-ADK or pBAD-DHFRs were grown in the presence of 100μg/ml of Ampicillin and 0.05% arabinose till an OD of 0.5 (approx. 3.5 hours). The cells were then harvested by centrifugation at 4C, and washed three times with 1×M9 salts in water. Approximately 25mg of cells were mixed with 300μl of 80:20 ratio of methanol:water that had been pre-chilled on dry ice. The cell suspension was immediately frozen in liquid nitrogen followed by a brief thawing (for 30 seconds) in a water bath maintained at 25C and centrifugation at 4C at maximum speed for 10 minutes. The supernatant was collected and stored on dry ice. This process of extraction of metabolite was repeated two more times. The final 900μl extract was spun down one more time and the supernatant was stored in −80C till used for mass spectrometry.

### *In vivo* cross-linking, immunoprecipitation (IP) and detection by mass-spec

BW27783 cells transformed with pBAD-empty, pBAD-ADK or pBAD-DHFRs were grown in the presence of 100μg/ml of Ampicillin and 0.05% arabinose for 4 hours. The cells were harvested by centrifugation at 4C, washed twice with 1×PBS and resuspended to a final OD of 5.0. For *in vivo* cross-linking, DSP (dithiobis(succinimidyl propionate)) was added to the cells to a final concentration of 2.5mM (stock 250mM in DMSO), and the reaction was allowed to proceed for 15 minutes at room temperature in a rotator/mixer. Extra cross-linker was quenched by adding 50mM Tris, pH 8.0 and kept for additional 30 minutes at room temperature in the rotator. The cells were collected by centrifugation, the final weight was determined, and cells were stored at −20C. For Co-IP, Invitrogen’s Dynabeads^®^ Co-Immunoprecipitation Kit was used with some small modifications.

For EcDHFR, pull-down was carried out using both anti-DHFR and anti-his antibodies in separate experiments. For DHFR 11, only anti-his antibodies were used. For ADK and Ras, polyclonal anti-ADK and anti-Ras antibodies were used.

### Activity measurement of the purified PurH

For the AICAR transformylase activity, ^10^Nformyl-THF was prepared from (6R,S)-5-formyl-THF (Schircks Laboratories, Jona, Switzerland) essentially as described in (Xu et al., 2004). For the assay, 250nM of PurH was incubated alone or with 0.1, 0.5, 2.5, 12.5 and 60μM of DHFRs or ADK for 30 minutes at room temperature in buffer (50mM Tis-HCl pH 7.4, 50mM β-ME and 25mM KCl) containing varying amounts of ^10^Nf-THF in a total volume of 95ul. The reaction was initiated by adding 5μl of 10mM AICAR (final concentration 0.5mM) and the increase in absorbance at 298nm was monitored.

### Activity assay for GlyA

20μM of GlyA were pre-incubated with varying concentrations of DHFRs/ADK (zero to 150μM) before addition of 0.2M glycine and 200μM 5f-THF (Schirch et al., 1985) and the amount of the ternary complex formed was monitored at 500nm. To determine of K_D_ for 5f-THF in the presence of added DHFR/ADK, the concentration of the latter was fixed at 60μM. The procedure followed is essentially described in (Fu et al., 2003).

## Supplementary Materials & Methods

### Strains, media, and growth conditions

Strains used for down-regulation of DHFR were derivatives of MG1655, while expression of pBAD plasmid in all cases were done in BW27783 cells (CGSC #12119) (Khlebnikov et al., 2001). Standard growth was conducted under the following conditions. Cells were grown from a single colony overnight at 37°C in M9 minimal salts supplemented with 0.2% glucose, 1mM MgSO4, 0.1% casamino acids, and 0.5 μg/ml thiamine (supplemented M9 medium). Overnight cultures were diluted 1/100 and grown at 37°C. Growth rate measurements were conducted for 12 hours in Bioscreen C system (Growth Curves USA). OD data were collected at 600nm at 15 min intervals. The resulting growth curves were fit to a bacterial growth model to obtain growth rate parameters (Zwietering et al., 1990).

### DHFR down-regulation and up-regulation

Generation of the strain with chromosomal controllable *folA* expression was previously described (Bershtein et al., 2013). Briefly, the operator sequence of the *lac* operon (lacO) was introduced upstream to −33 and −10 promoter signals of the chromosomal *folA* gene in a strain carrying Z1 cassette with *lacI* gene under constitutive promoter 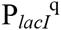(Lutz and Bujard, 1997). Under the saturated IPTG concentration (0.6 mM), the resulted strain produced 20–25% of the basal DHFR expression level found in wild-type MG1655 strain, as determined by Western Blot (see below). Decrease in the IPTG concentration resulted in further drop of DHFR production. Design of the strains carrying orthologous DHFR replacements was previously described (Bershtein et al., 2015).

For over-expression, BW27783 cells transformed with pBAD plasmid were subjected to varying arabinose concentrations (from 0 to 0.05%) and intracellular abundance was calculated by Western Blot. For metabolite complementation, overnight culture (grown without any metabolite) was diluted 1/100 in supplemented M9 medium containing 1mM of different metabolites.

### Intracellular protein abundance

Cells were grown in supplemented M9 medium for 4 hours at 37°C, chilled on ice for 30 min and lysed with 1×BugBuster (Novagen) and 25units/ml of Benzonase. Intracellular DHFR, ADK and ras amounts in the soluble fraction were determined by SDS-PAGE followed by Western Blot using rabbit anti-DHFR (E.coli)/anti-ADK/anti-ras

**Figure S1 (Related to Fig.1):**
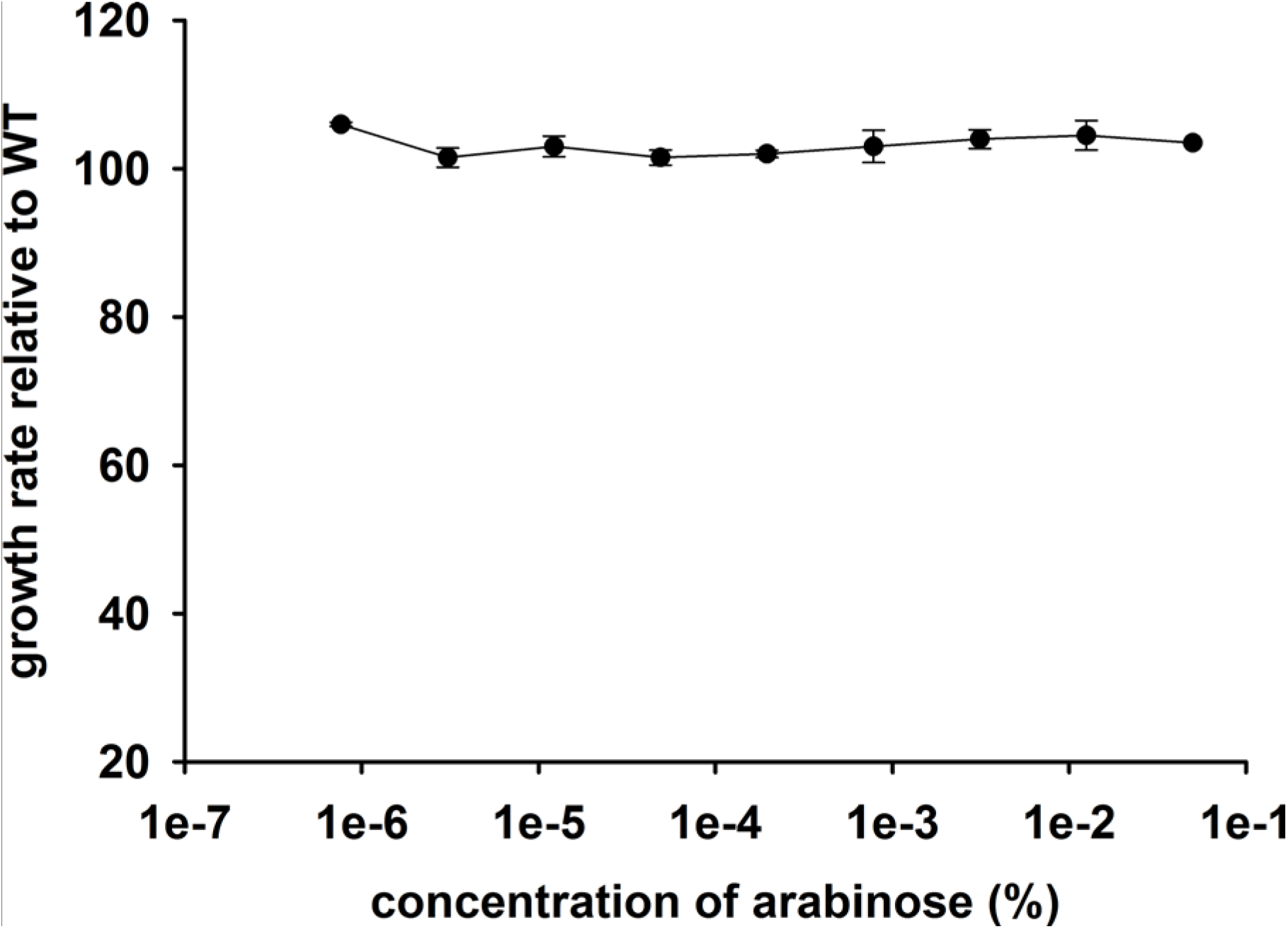
Growth rate (relative to untransformed cells) of *E.coli* cells transformed with an empty pBAD plasmid over a range of arabinose concentrations. The empty plasmid does not show any toxicity.

**Figure S2 (Related to Fig.1):**
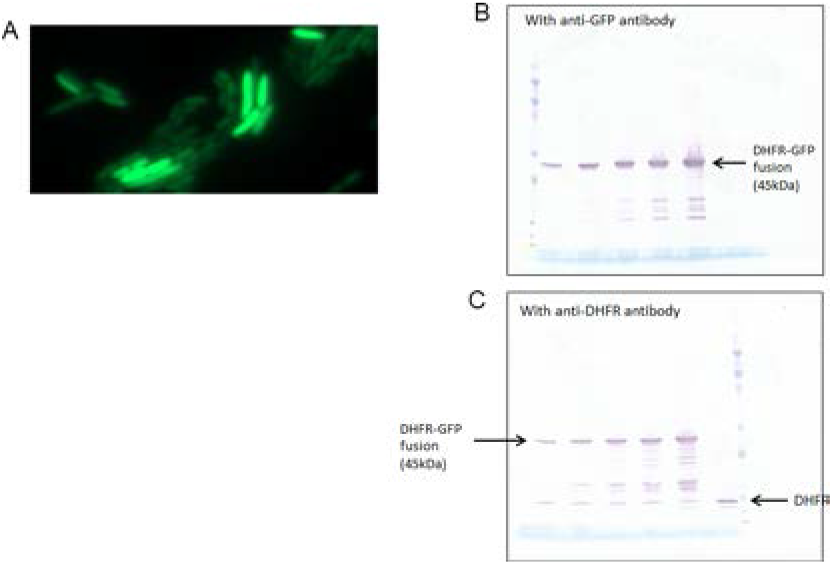
Over-expressed DHFR does not aggregate inside the cell. (A) DHFR fused to monomeric superfolder GFP was expressed from pBAD plasmid, and the fluorescence was detected. The fluorescence was uniformly spread over the cells, ruling out any substantial aggregation. (B) Over-expressed fusion protein is not cleaved inside the cell, as both anti-DHFR and anti-GFP antibodies showed very minor fraction of cleaved proteins.

**Figure S3 (Related to Fig.2):**
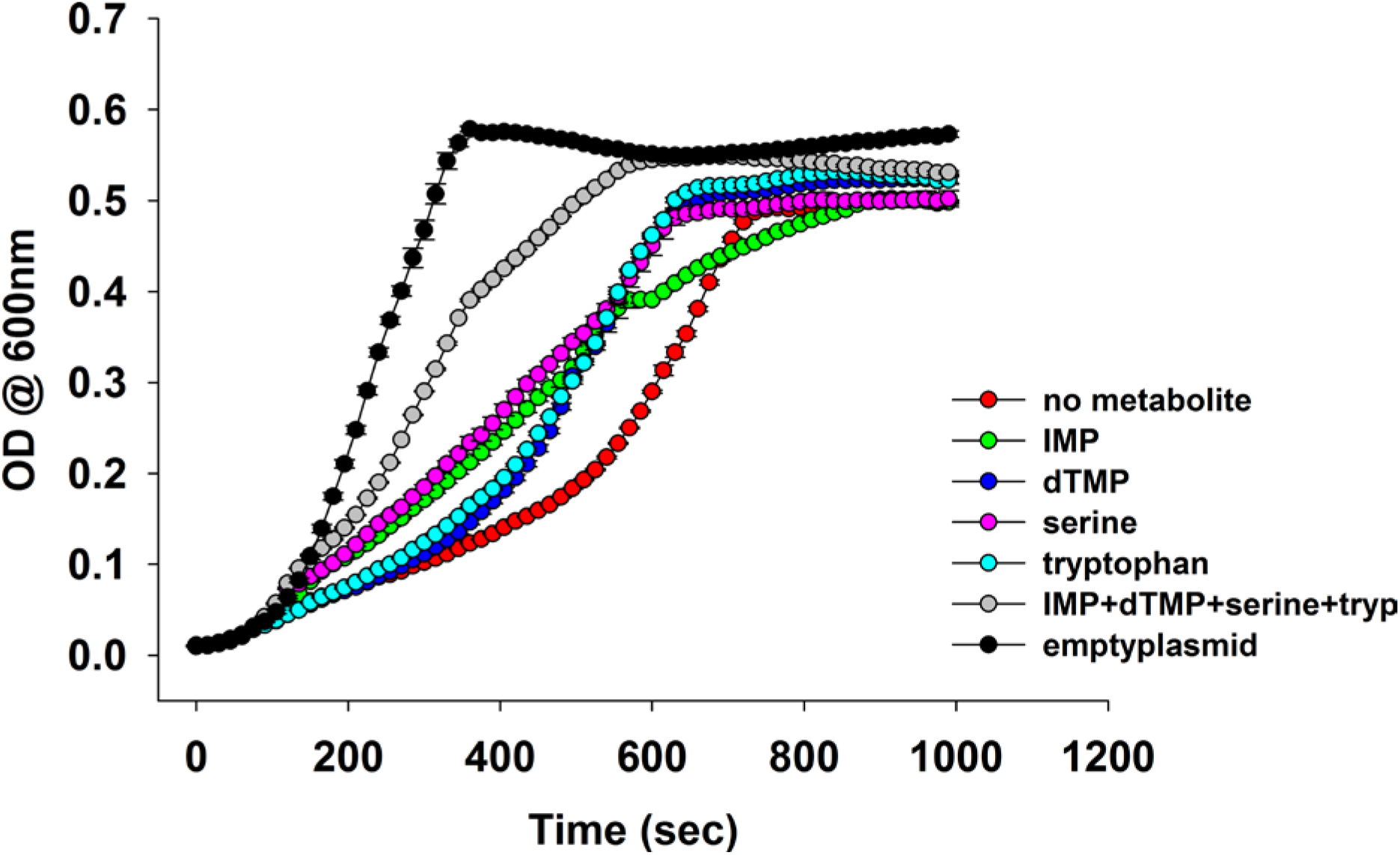
Overlay of growth curves of *E.coli* over-expressing DHFR and those in the presence of various metabolites. Addition of 1mM IMP, dTMP, serine and tryptophan to the medium showed a marked improvement in the growth.

**Figure S4 (related to Fig.3):**
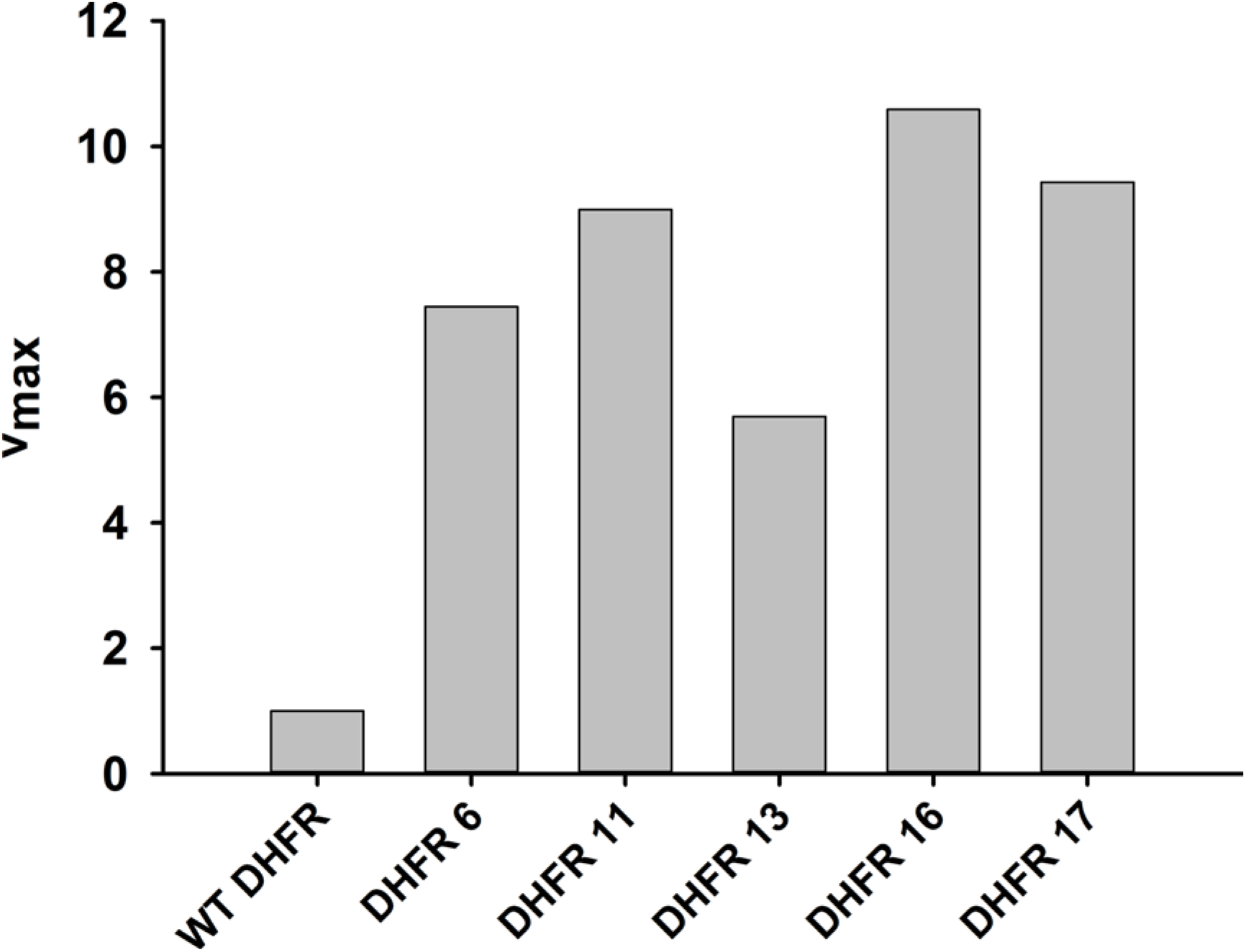
Activity (v_max_) of over-expressed DHFR from whole cell lysate. The initial rate of conversion of 50μM Dihydrofolate (DHF) to THF was measured in the presence of 100μM NADPH in 50mM MES buffer for *E.coli* over-expressing different DHFRs. As expected from their *in vitro* activity parameters, over-expressed orthologous DHFR proteins were more active than *E.coli* DHFR even in the lysate.

**Figure S5 (Related to Fig.5):**
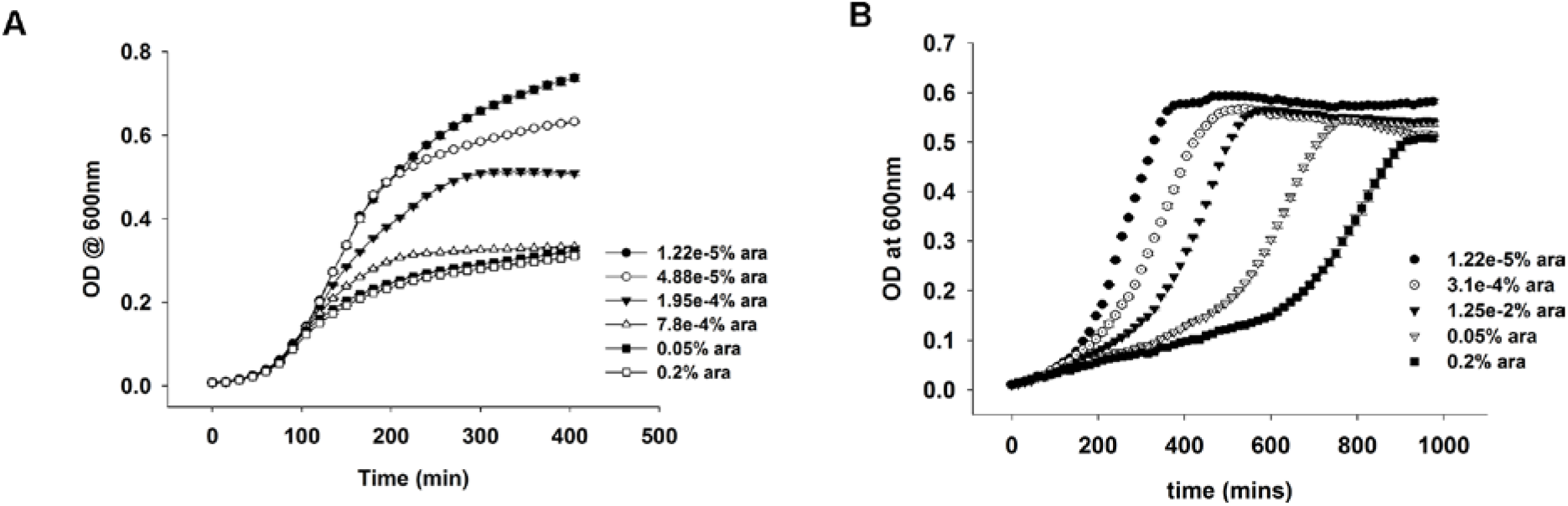
OD vs time curves for DHFR over-expression in (A) rich media (LB) and (B) in M9 medium supplemented with amino acids. In LB, the growth rate is not affected due to over-expression, though the cells cannot grow to the same saturation ODs. In M9 however, the growth rate is severely affected, while the saturation ODs remain largely unchanged.

**Figure S6 (Related to Fig.5):**
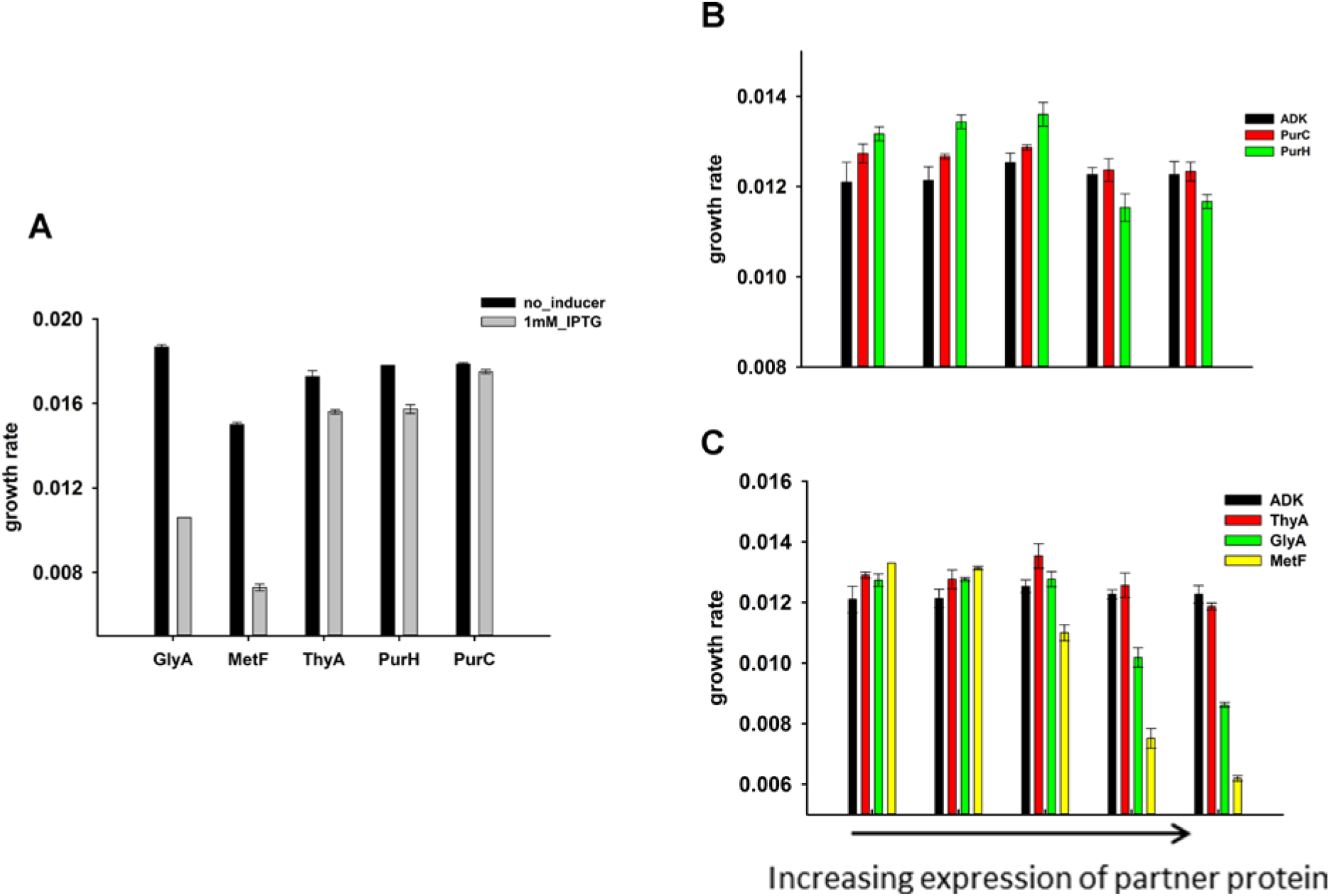
Effect of over-expression of purine biosynthesis and 1-carbon metabolism pathway proteins on growth rates of *E.coli* (A) alone and (B) on the background of DHFR over-expression. GlyA and metF are significantly toxic at the highest induction level, ThyA and PurH show moderate toxicity while PurC was found to be non-toxic. On the background of DHFR over-expression (basal expression from pTRC plasmid), expression of the partner proteins was beneficial at low expression levels, however eventually became toxic at higher inducer concentrations.

**Figure S7 (Related to Fig.6):**
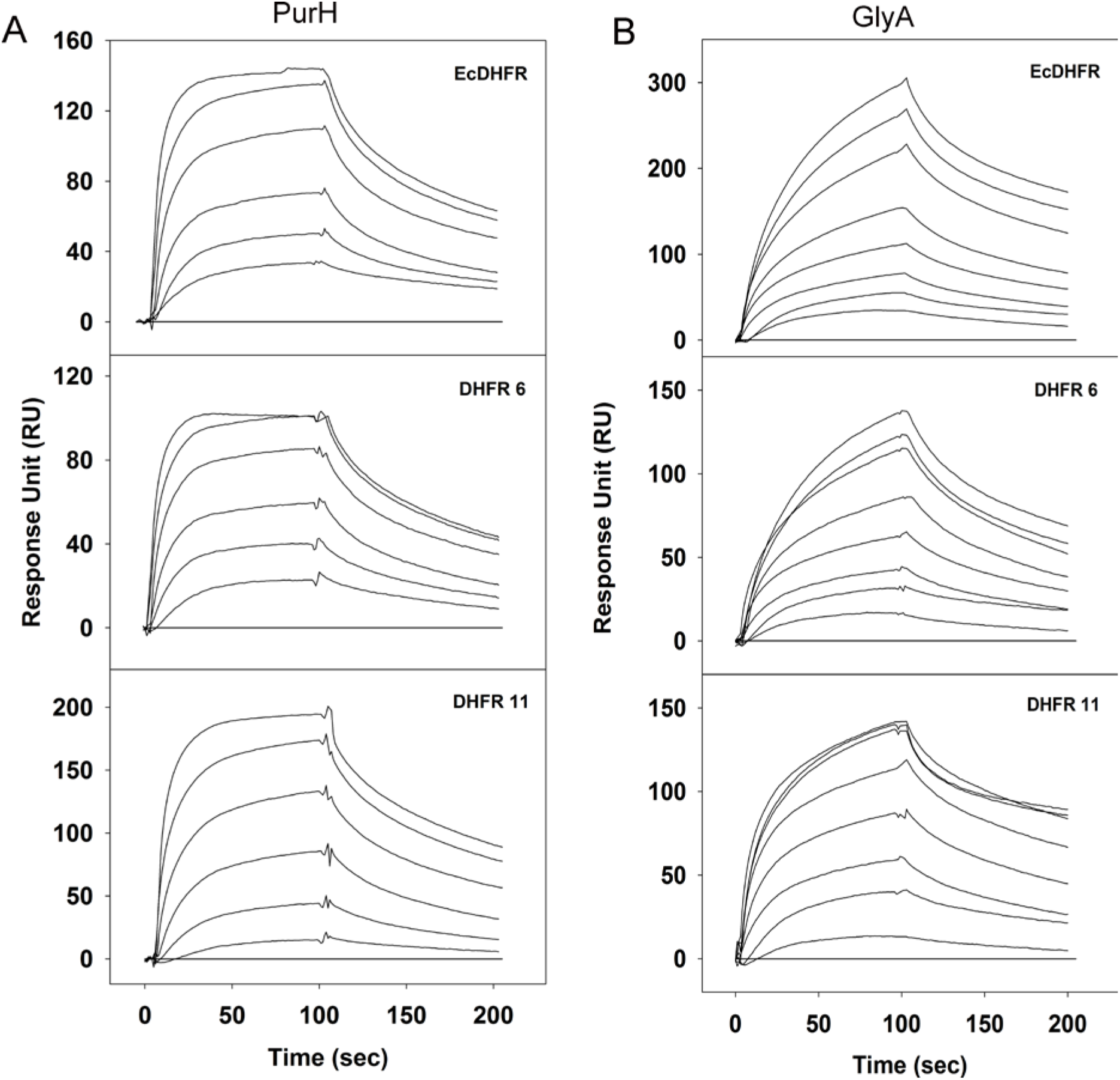
Binding of purified PurH and GlyA proteins with EcDHFR, DHFR 6 and 11 *in vitro* detected using surface plasmon resonance. 700-900 RUs of purified his-tagged DHFRs were immobilized on a CM5 chip and (A) PurH and (B) GlyA proteins were flown over the surface at 25°C. For panel (A), curves from bottom to top represent 1.5μM, 3μM, 6μM, 12μM, 24μM and 48μM of purified PurH. For panel (B), curves from bottom to top represent 1μM, 2μM, 4μM, 8μM, 16μM, 32μM, 48μM and 64μM of purified GlyA. His-tagged ADK protein was immobilized on the negative control channel and was subtracted from all binding data (baseline shown in all three plots).

**Figure S8 (related to Fig.6):**
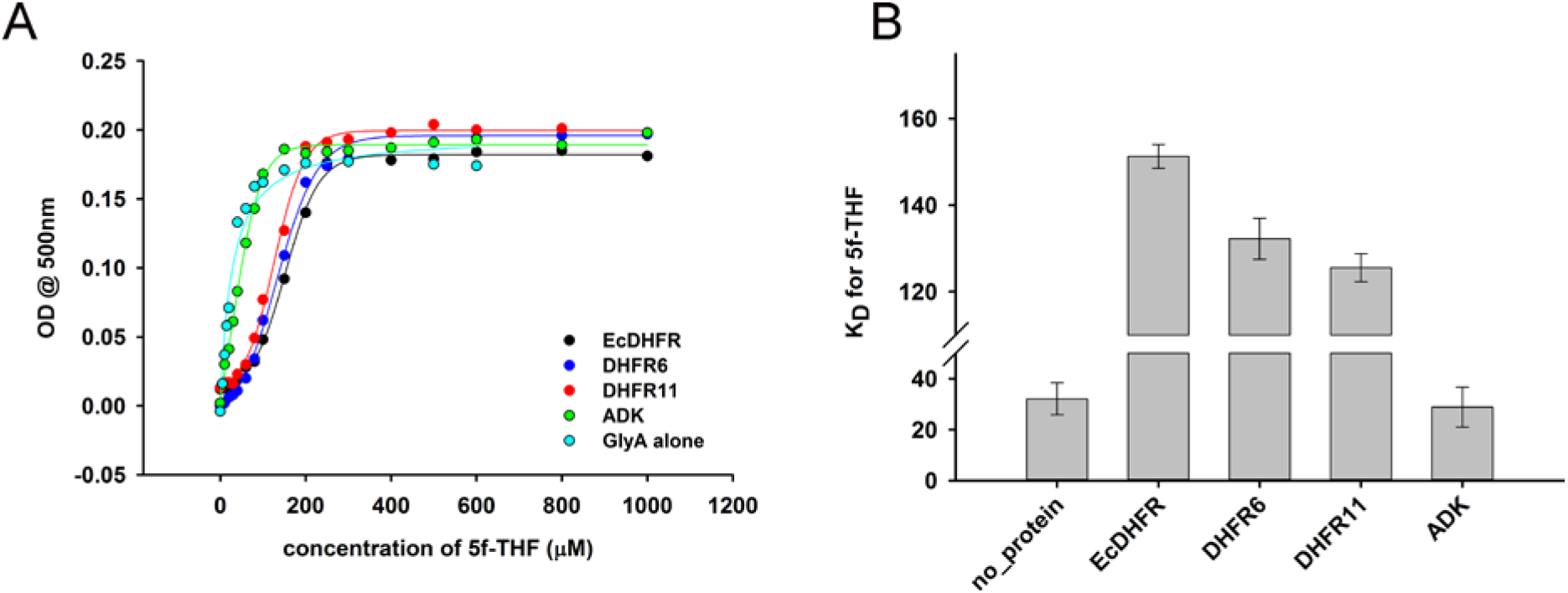
Determination of equilibrium dissociation constant (K_D_) of binding of 5f-THF to GlyA (A) Formation of a ternary complex (GlyA-PLP+glycine+5f-THF) was monitored at 500nm as a function of increasing amounts of 5f-THF in the presence of different DHFRs/ADK. 20μM GlyA was pre-incubated with 60μM of DHFRs/ADK and the concentration of 5f-THF was varied from 0 to 1mM. (B) While addition of both EcDHFR as well as the orthologs increased the K_D_ of 5f-THF binding to GlyA substantially, EcDHFR caused the largest increase, and therefore inhibited GlyA the most. Presence of ADK did not perturb the K_D_, indicating that the effect of DHFRs was highly specific.

**Figure S9 (related to Fig.6):**
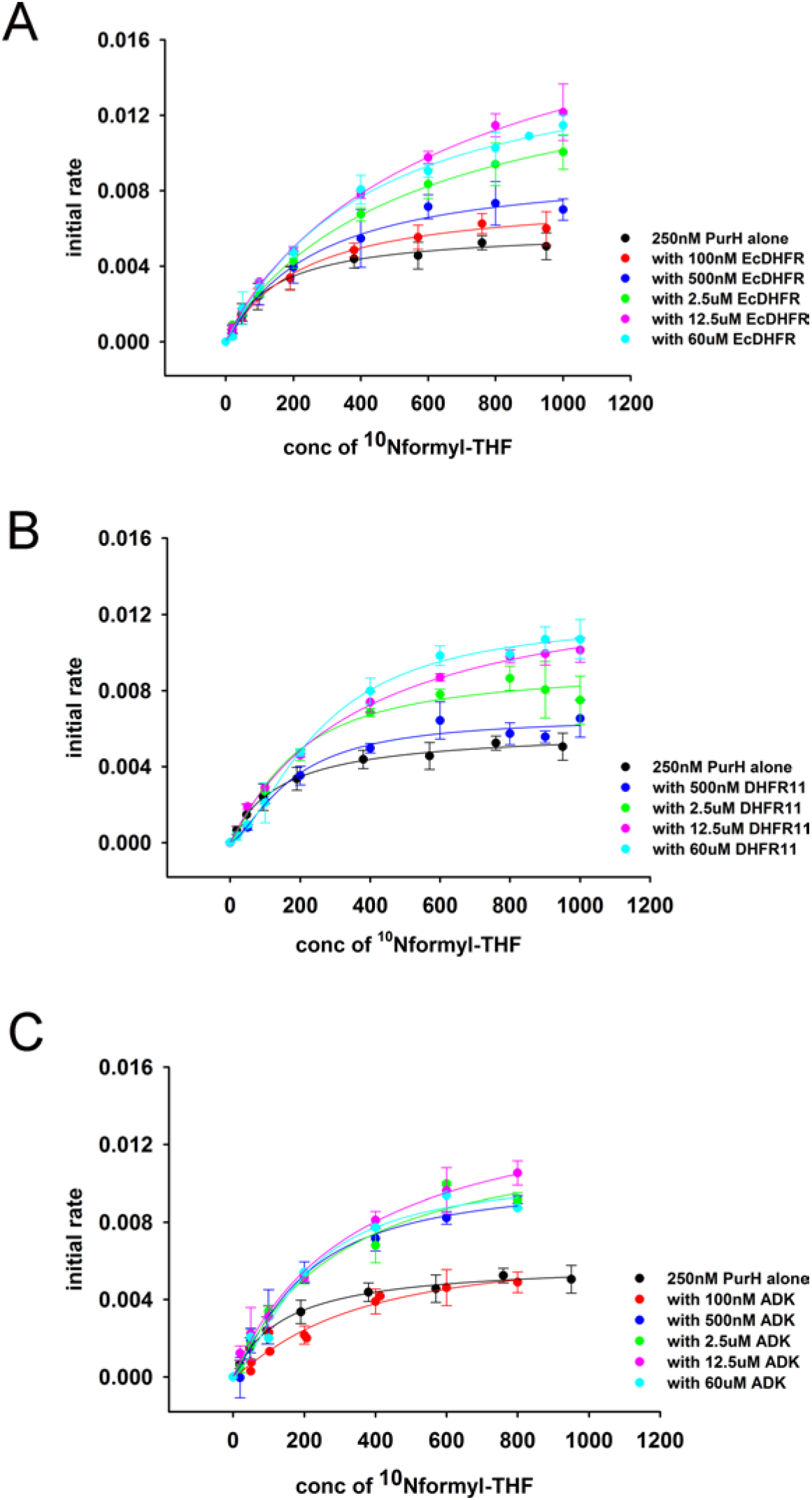
Overlay of initial rate (maximum rate) of PurH activity as a function of ^10^Nf-THF concentration at different concentrations of added protein (A) EcDHFR (B) DHFR 11 and (C) ADK. For all proteins, there is steady increase in the initial rates with addition of extra protein.

**Table S1:** Metabolomics data

| Table S1: Metabolomics data |  |  |  |  |  |  |  |  |
| --- | --- | --- | --- | --- | --- | --- | --- | --- |
|  |  | AICAR | dUMP | dTMP | dTTP | AMP | GMP | IMP |
|  | empty plasmid | 0 | 0 | 1.00 | 1.00 | 1.00 | 1.00 | 1.00 |
|  | ADK | 0 | 0 | 1.51 | 1.47 | 1.33 | 0.92 | 0.95 |
|  | E.coli DHFR | 35217676 | 122171 | 0.13 | 0.00 | 0.26 | 0.69 | 0.25 |
|  | DHFR 6 | 0 | 6399 | 1.04 | 1.43 | 1.06 | 1.04 | 0.55 |
|  | DHFR 11 | 0 | 4387 | 0.88 | 1.21 | 0.81 | 1.08 | 0.63 |

**Table S2:**
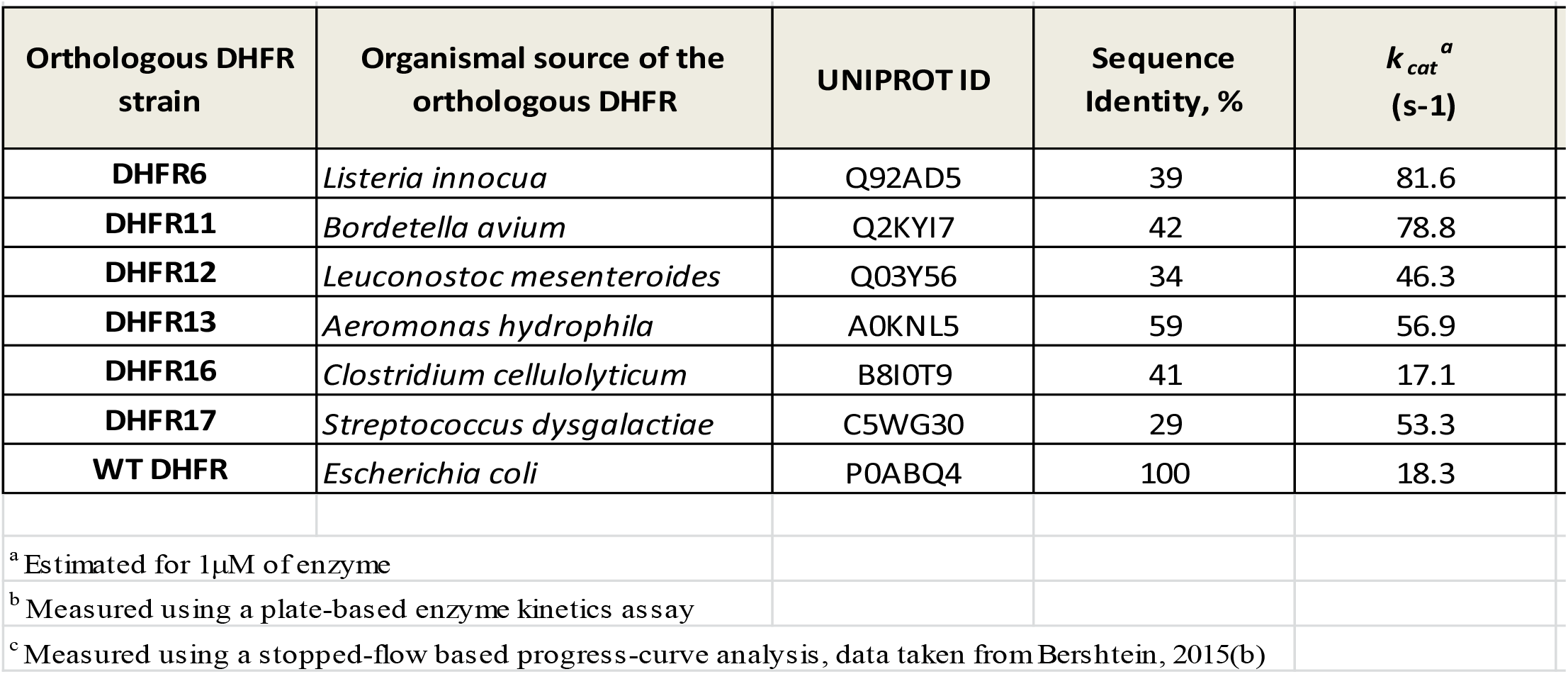
Molecular properties of orthologous DHFR proteins

**Tables S3 and S4 are available upon request**

**Table S5:** Kinetic parameters for binding of purified purH and glyA proteins to su

| | | | $k_{\text{on}}$ ($\text{M}^{-1} \text{s}^{-1}$) | $k_{\text{off}}$ ($\text{s}^{-1}$) | $K_{\text{D}}$ (M) |
| --- | --- | --- | --- | --- | --- |
|  | PurH | EcDHFR | 9.10E+03 | 2.50E-02 | 2.75E-06 |
|  |  | DHFR 6 | 8.00E+03 | 2.36E-02 | 3.00E-06 |
|  |  | DHFR 11 | 6.40E+03 | 2.10E-02 | 3.30E-06 |
|  | GlyA | EcDHFR | 2.45E+03 | 1.96E-02 | 8.00E-06 |
|  |  | DHFR 6 | 3.90E+03 | 2.05E-02 | 5.30E-06 |
|  |  | DHFR 11 | 5.90E+03 | 2.36E-02 | 4.00E-06 |

